# Predicting Individual Cell Division Events from Single-Cell ERK and Akt Dynamics

**DOI:** 10.1101/2021.09.14.460349

**Authors:** Alan D Stern, Gregory R Smith, Luis C Santos, Deepraj Sarmah, Xiang Zhang, Xiaoming Lu, Federico Iuricich, Gaurav Pandey, Ravi Iyengar, Marc R Birtwistle

## Abstract

Predictive determinants of stochastic single-cell fates have been elusive, even for the well-studied mammalian cell cycle. What drives proliferation decisions of single cells at any given time? We monitored single-cell dynamics of the ERK and Akt pathways, critical cell cycle progression hubs and anti-cancer drug targets, and paired them to division events in the same single cells using the non-transformed MCF10A epithelial line. Following growth factor treatment, in cells that divide both ERK and Akt activities are significantly higher within the S-G2 time window (∼8.5-40 hours). Such differences were much smaller in the pre-S-phase, restriction point window which is traditionally associated with ERK and Akt activity dependence, suggesting unappreciated roles for ERK and Akt in S through G2. Machine learning algorithms show that simple metrics of central tendency in this time window are most predictive for subsequent cell division; median ERK and Akt activities classify individual division events with an AUC=0.76. Surprisingly, ERK dynamics alone predict division in individual cells with an AUC=0.74, suggesting Akt activity dynamics contribute little to the decision driving cell division in this context. We also find that ERK and Akt activities are less correlated with each other in cells that divide. Network reconstruction experiments demonstrated that this correlation behavior was likely not due to crosstalk, as ERK and Akt do not interact in this context, in contrast to other transformed cell types. Overall, our findings support roles for ERK and Akt activity throughout the cell cycle as opposed to just before the restriction point, and suggest ERK activity dynamics are substantially more important than Akt activity dynamics for driving cell division in this non-transformed context. Single cell imaging along with machine learning algorithms provide a better basis to understand cell cycle progression on the single cell level.

## Introduction

The mammalian cell cycle is in large part driven by growth factor activation of the Ras-ERK^1–5^ and the PI3K-Akt^2,6–11^ pathways. Growth factors cause auto-phosphorylation of receptor tyrosine kinases (RTKs). For the ERK pathway, RTK phosphorylation recruits the guanine exchange factor SOS to the membrane, catalyzing the exchange of GDP for GTP bound to Ras, initiating Raf activation ^6,12–16^. This in turn activates the MEK-ERK phosphorylation cascade. When activated, the effector kinases ERK1/2 translocate from the cytoplasm to the nucleus and activate transcriptional regulators such as Elk-1 and CREB^17,18^ ^19^. These transcriptional regulators induce immediate early genes (IEGs) like *c-fos*^17,18^ that then contribute to the expression of cyclin D1 ^4,6,8,19–21^, a key step in S-phase entry^22^.

RTK activation can also initiate Akt pathway signaling. RTK autophosphorylation recruits adapter proteins like insulin receptor substrate (IRS-1) and GRB2-associated binder (GAB) ^23–26^. These proteins in turn recruit Phosphatidylinositol (PtdIns) 3-kinase (PI3K) to the membrane ^21,26–28^ where it phosphorylates the membrane phospholipid PtdIns (4,5) P2 (PIP2), generating PtdIns (3,4,5) P3 (PIP3). PIP3 recruits pleckstrin homology domain (PH)-containing proteins to the membrane such as phosphatidylinositol-dependent kinase-1 (PDK1)^29^ and the serine/threonine protein kinases Akt1/2 ^21,30^. PDK1 phosphorylates Akt’s activation loop followed by mTORC2 phosphorylation of a second site on Akt for full activation ^6,7,21^. This doubly phosphorylated, activated Akt promotes cell cycle progression by: (i) promoting protein translation via 4E-BP and p70S6K ^6,21^, (ii) promoting cyclin D1 ^20,31^ CDK4/6, c-Myc, and E2F activity ^32^ and (iii) inhibiting p21 and p27 ^33^ (cyclin-dependent kinase inhibitors).

While ERK and Akt pathways have established roles prior to the restriction point marked by S-phase entry, the extent to which they are informative of cell cycle completion after S-phase is less clear. Beyond S-phase, studies suggest that Ras-ERK ^34–41^ and PI3K-Akt ^30,42,43^ may contribute towards regulating G2 progression. ERK activity was shown to play a role in the duration of DNA damage-induced G2 arrest^44^. Transient ERK activity maintains G2 arrest, whereas sustained ERK activity promotes escape by reducing p53 levels, and inducing the expression of pro-mitotic factors such as Plk1 and cyclin B ^41^. Akt activity also contributes to G2-M progression as its inhibition is associated with reduced cyclin B levels, promoting Chk1 activity and G2 arrest ^42^. These observations motivate a closer look at determining how ERK and Akt dynamics are informative of cell cycle completion after the canonical restriction point.

On a single cell level, both ERK and Akt activity dynamics have substantial cell-to-cell and dynamic variation, exhibiting complex pulses and more simple steady activity ^1,27,45–53^. Such variation, when coupled with the observations that cell cycle progression is also heterogeneous ^1,54,55^, have prompted investigations into the correlation between dynamics and cell cycle fate in single cells. What determines proliferation on a single cell level? What relative contributions do ERK and Akt activity have to the decision of individual cells to divide? Much prior work has focused on activity dynamics. Both Ras-ERK ^56–58^ and PI3K-Akt ^59^ exhibit biphasic growth factor-induced activation dynamics, with a transient peak followed by sustained activity hours later. The dynamics of each phase contributes differently towards driving progression to S-phase and is cell type dependent ^59–61^. Live-cell imaging and analysis of recently divided sister cells reveal that time-integrated ERK activity has some predictive power of the timing to S-phase entry^1^. Time-integrated ERK dynamics were found to influence proliferation decisions in daughter cells ^50^. Predicting PC12 cell differentiation/proliferation decisions required both ERK and Akt activity dynamics to best define the decision boundary between these two cell fate outcomes ^62^. Yet, the extent to which both ERK and Akt activities throughout the cell cycle are predictive of division in single cells remains unclear.

Here, we use live-cell imaging to pair measurements of growth factor-induced ERK and Akt activity to cell division outcomes in the same single cells. We aim to assess the extent to which these activities are associated with cell cycle progression beyond S-phase entry, and to evaluate their ability to predict single cell division responses jointly in the well-established non-transformed breast epithelial MCF10A cell line, a model system that is commonly used to study epithelial signaling biology and cell division control ^26,63–68^. We found that following treatment of synchronized cells with growth factors EGF and insulin, both ERK and Akt activity are significantly higher within the S-G2 interval in dividing cells. Such differences were much smaller in the pre-S-phase window, which is traditionally associated with ERK and Akt activity dependence^59–61^, suggesting unappreciated roles for ERK and Akt in S through G2. These higher activities could classify division events with AUC=0.76. Surprisingly, ERK activity dynamics alone enable AUC=0.74, suggesting Akt activity dynamics contribute little to the decision governing cell division in this context. Interestingly, we found that ERK and Akt activities are less correlated in cells that divide. Network reconstruction experiments demonstrated that this correlation behavior was not due to crosstalk, as ERK and Akt do not interact in this context, in contrast to other cell types ^69^. Overall, our findings support roles for ERK and Akt activity throughout the cell cycle as opposed to just before the restriction point, and suggest ERK activity dynamics are substantially more important than Akt activity dynamics for driving cell division in this non-transformed context.

## Results

### Predictability of Cell Division Events from Univariate ERK and Akt Dynamics

To evaluate if ERK or Akt signaling dynamics are predictive of cell division we first conducted a series of live cell imaging experiments in MCF10A cells that express either ERK^46,70^ or Akt^48^ kinase translocation reporters (KTRs), but not yet both at once, and paired those single cell dynamics to cell division events (Fig. 1A). We first verified that cell cycle progression and division are related to ERK and Akt activity dynamics in MCF10A cells using small molecule inhibitor experiments (Fig S1). KTR-expressing cells were G0-synchronized by serum and growth factor starvation for 24 hours. After acquiring 1 hour of baseline ERK or Akt activity, cells were treated with EGF and insulin, growth factors that promote cell division in MCF10A cells^71^. Images were acquired every 15 minutes for 48 hours, and then single-cell data for kinase activity and division outcome were extracted using custom-built image processing pipelines (see Methods). Dynamic regimes of KTR specificity were determined using two independent (four total) MEK and Akt inhibitors (Fig. S2). ERK KTR was found to be specific in all regimes explored here, whereas the Akt KTR was found to be specific >∼ 1 hour after EGF and insulin co-stimulation.

**Figure 1.**
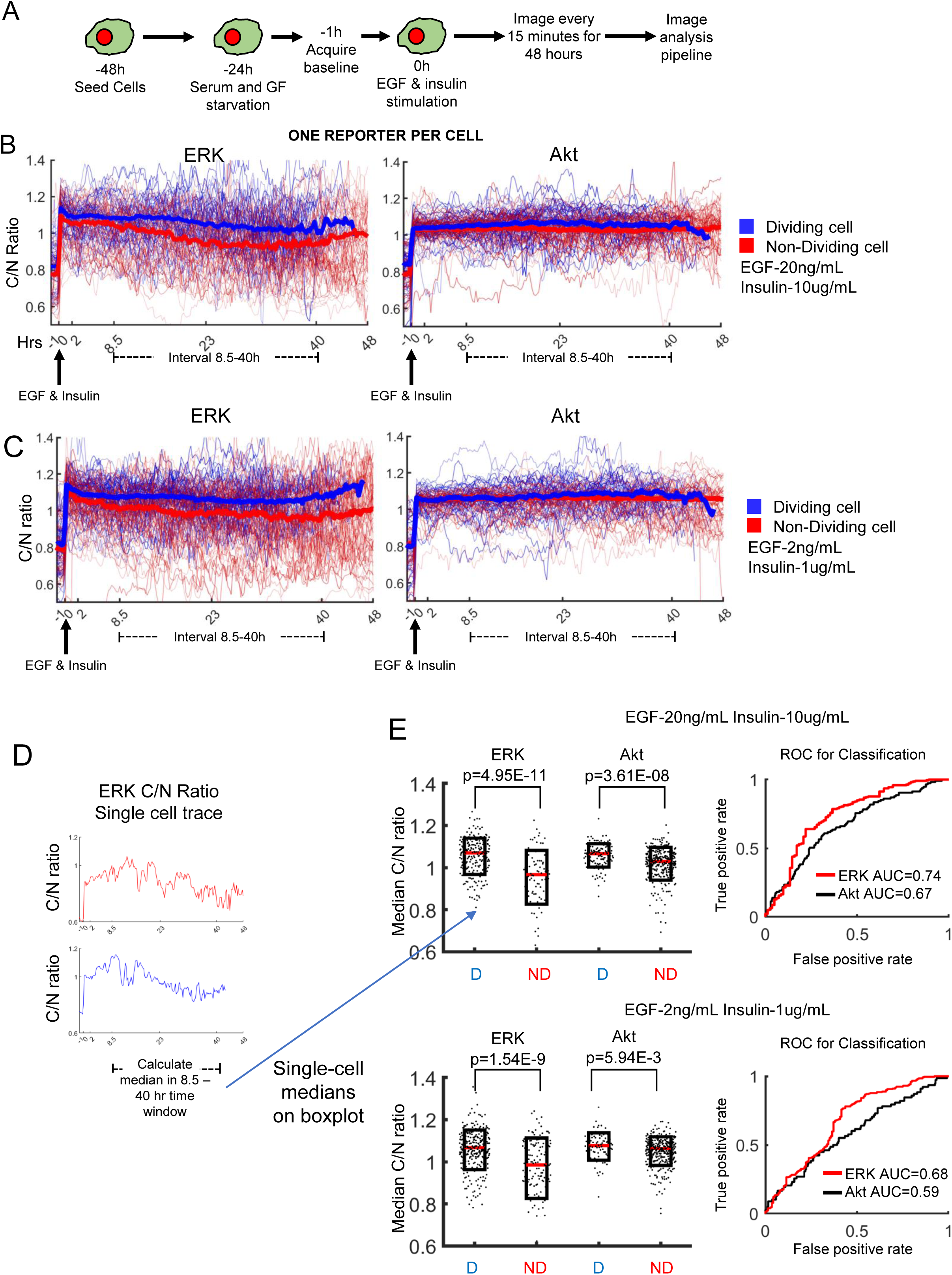
Evaluating the predictability of cell division from univariate single cell ERK or Akt dynamics. In these experiments, cells were either expressing the ERK or the Akt KTR. (**A**) Cell treatment workflow for pairing single cell KTR dynamics to cell division. ERK or Akt KTR expressing MCF10A cells were seeded, allowed to attach overnight, and then serum and growth factor starved. Following starvation, baseline images were acquired, cells were treated with EGF and insulin, and then imaged every 15 minutes for 48 hours. Images were quantified using the analysis pipeline described in the methods. (**B, C**) Quantified ERK or Akt KTR dynamics paired to division events for EGF and Insulin doses that match those used in culture medium (**B**) or 10-fold less (**C**). Single cell traces of dividing (blue) and non-dividing (red) cells are shown with thin lines, and population median (per time point) is shown with thick lines. (**D**) Left, representative single cell trace of ERK KTR for a dividing (blue) or non-dividing (red) cell. Median ERK activity within the 8.5-40 hour interval for each cell becomes a single dot in the boxplots. (**E**) Left: notBoxplots for single cell median ERK or Akt activity within the 8.5-40 hour interval for EGF and Insulin doses that match those used in culture medium (top) or 10-fold less (bottom). p-values for right tailed rank-sum test were calculated at the 95% confidence interval. D: dividing; ND: Non-dividing. Right: ROC curve for classifying cell division events from single cell median ERK (red) or Akt (black) activity using a logistic regression model.

Single cell traces of ERK or Akt activity (thin lines) along with the population median (bold line) show rapid activation following growth factor treatment, which largely persists for the duration of the experiment, without recognizable pulsing (Fig. 1B,C). In (blue) dividing cells, population median ERK and Akt activity dynamics are higher throughout the cell cycle compared to non-dividing cells, with larger differences evident for ERK. In the pre-S-phase entry window (∼< 8 hours after growth factor treatment), there are slight differences between dividing and non-dividing cells in terms of population median ERK and Akt dynamics. These differences grow larger in the subsequent 8.5—40 hour interval post growth factor addition, which largely corresponds to S and G2 phases. These trends were also evident with 10-fold less concentration of growth factors (Fig. 1C). These results suggest that ERK and Akt activity may have importance after S through G2 phase.

To assess the statistical significance of this finding, we calculated the median ERK or Akt activity for individual single cells within the 8.5 – 40 hour window post-growth factor treatment, and then compared median activity between dividing and non-dividing cells with the rank-sum test (Fig. 1D-E). Individual dots in the boxplot represent the median ERK or Akt activity calculated within the 8.5 – 40 hour interval in a single cell. These median single cell activities were significantly different in dividing vs. non-dividing populations (Fig. 1E). Yet, there is substantial overlap in the two populations. We evaluated whether single cell median ERK or Akt dynamics are predictive of cell division using a logistic regression classification model, and ROC analysis to quantify the outcome. Both ERK and Akt dynamics have some predictive power for cell division under high and low growth factor conditions, with high growth factor conditions having slightly elevated predictive power, as quantified by the area under the ROC curve (AUC). ERK dynamics have more predictive power than Akt dynamics. Yet, the best achieved AUC is 0.74, indicating there are other factors driving differences in cell division fate.

### Predictability of Cell Division Events Using Measurements of Both ERK and Akt Dynamics in the Same Single Cells

As shown above, ERK and Akt activity dynamics alone contain information about subsequent cell division. Would simultaneous measurements of both ERK and Akt activity dynamics in the same single cell improve cell division predictions? To answer this question, we performed a similar experiment as described above using dual reporter expressing MCF10A cells (see Methods). For the duration of the time course, population median ERK and Akt dynamics are again elevated in dividing cells compared to non-dividing cells (Fig. 2A), with larger differences observed in the 8.5 – 40 hr interval. To evaluate median ERK and Akt dynamics as bivariate predictors of cell division, we trained a support vector machine (SVM) classifier (Fig. 2B). ROC evaluation of SVM performance shows some, albeit small improvements from the ERK-only classifier performance (AUC = 0.76 vs. 0.74 and 0.72 vs. 0.68, Fig. 2D). These results were confirmed in an independent experiment (Fig. S3A-C). Thus, Akt dynamics add comparatively little new information to ERK dynamics for predicting single cell division events in this context.

**Figure 2.**
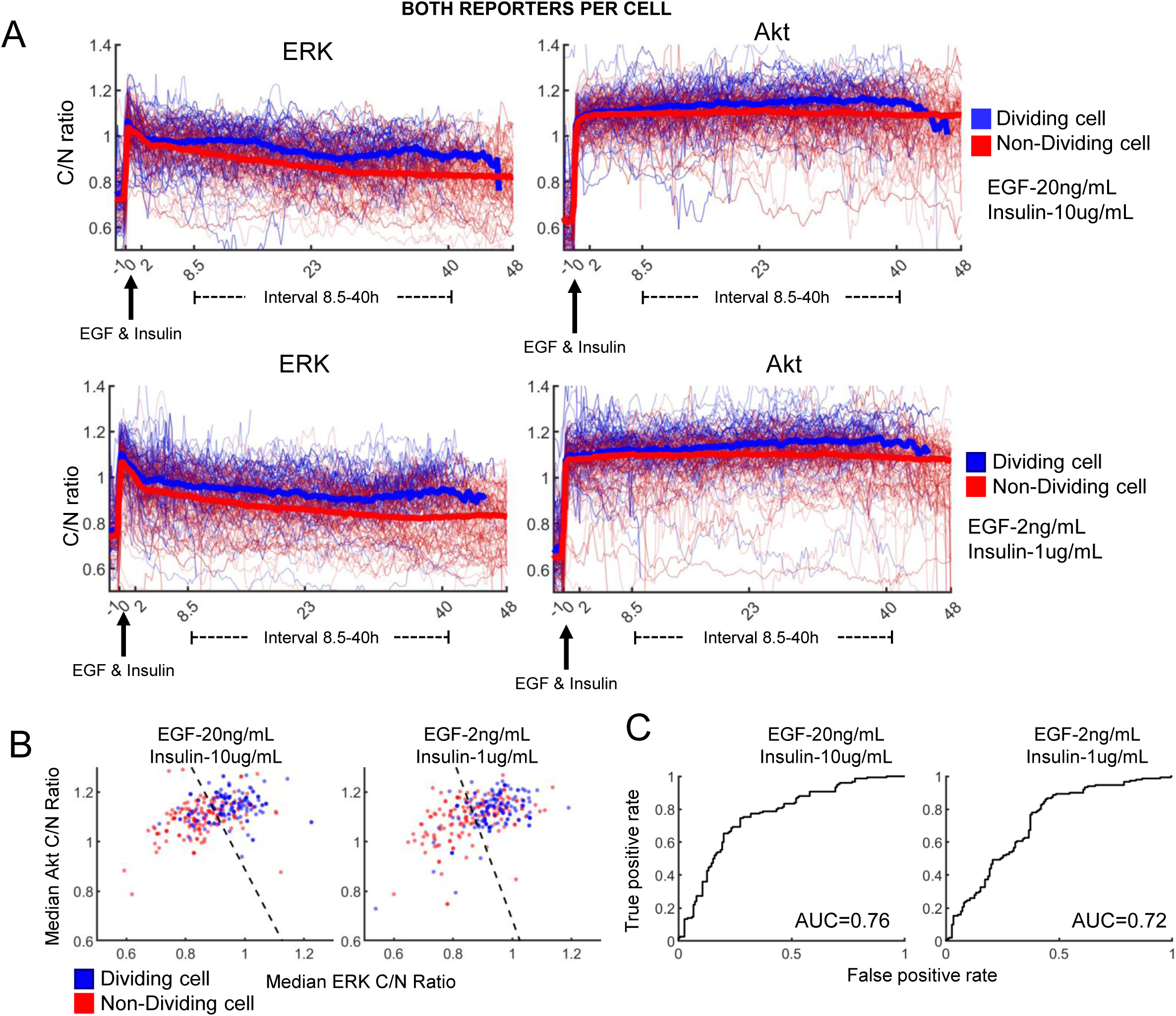
Evaluating the predictability of cell division from paired, bivariate single-cell ERK and Akt dynamics. In these experiments, cells were expressing both the ERK and Akt KTR simultaneously. (**A**) Quantified ERK and Akt dynamics for EGF and Insulin doses that match those used in culture medium (top) or 10-fold less (bottom). Single cell traces of dividing (blue) and non-dividing (red) cells are shown with thin lines, and population median (per time point) is shown with thick lines. (**B**) Scatter plot of ERK vs. Akt KTR median activity in the 8.5-40 hour window from 150 randomly sampled cells. Each dot is a single cell. Dividing cells are blue and non-dividing cells are red. The dotted line is the SVM hyperplane for classifying dividing and non-dividing cells. Left and right are high and low growth factor concentrations, respectively. (**C**) ROC curve for SVM classification performance for EGF and Insulin doses that match those used in culture medium (left) or 10-fold less (right).

### Measurements of Central Tendency Dominate Predictive Dynamic Features

The above analysis focused on simple median features of ERK and Akt dynamics as predictors of cell division. However, it was not clear *a priori* what dynamic features may be relevant to this prediction task. To determine if there are additional time series features that can improve cell division predictions, we utilized the machine learning-based highly comparative time-series analysis (hctsa) package ^72,73^. We calculated hctsa features on raw time series data for dividing and non-dividing cells in dual ERK and Akt KTR reporter expressing lines. Features that best discriminate dividing from non-dividing cells are all measures of central tendency (Table 1). Thus, we conclude that the above analysis focused on median activity is likely to be sufficient for assessing how predictive ERK or Akt dynamics are for cell division events in the studied context.

**Table 1.** The top 5 features of ERK and Akt signaling dynamics for prediction of cell division as determined by HCTSA analysis of dual ERK and Akt KTR expressing lines under 20ng/mL EGF, and 10ug/mL insulin. The top features shown are related to measurements of central tendency using different operation parameters corresponding to the values within the brackets []. These operations parameters and their corresponding functions can be found in hctsa documentation.

| ERK KTR Features |  | Akt KTR Features |  |
| --- | --- | --- | --- |
| Operation (HCTSA) | -log10 p-value (Rank sum) | Operation (HCTSA) | -log10 p-value (Rank sum) |
| [10] midhinge | 3.39 | [16] rms | 6.79 |
| [16] rms | 3.32 | [6] trimmed mean_5 | 6.74 |
| [5] trimmed mean_1 | 3.31 | [7] trimmed mean_10 | 6.72 |
| [2] mean | 3.30 | [8] trimmed mean_25 | 6.72 |
| [3] harmonic_mean | 3.29 | [5] trimmed mean_1 | 6.71 |

### Inferring the Topology of the ERK-Akt Network

Information content is related to correlation, so we investigated the extent to which ERK and Akt dynamics in the same single cells were correlated, looking across every cell and every time point (Fig. 3A, replicate Fig. S3D). Interestingly, in dividing cells, single cell ERK and Akt dynamics within the 8.5-40 hour window are significantly less correlated than in non-dividing cells, at both high and low growth factor doses. Network topology can strongly influence correlated behaviors. In different studies, ERK and Akt have been reported to exhibit very different network behavior, such as cross-pathway activation, inhibition^6,20,21,69,74–79^ and non-interaction^23,69,80^. Factors such as cell type and growth factor context can influence these discrepant network topologies^3,69^. Previous work conducted in panel of growth factors and cell lines show varying probabilities of forming an interaction network edge between ERK and Akt^69^. The differences in network edge formation can affect downstream signaling and cell fate decisions^3^. Could ERK and Akt network topology be dynamic, and give insight into the division-related correlated behaviors observed above?

**Figure 3.**
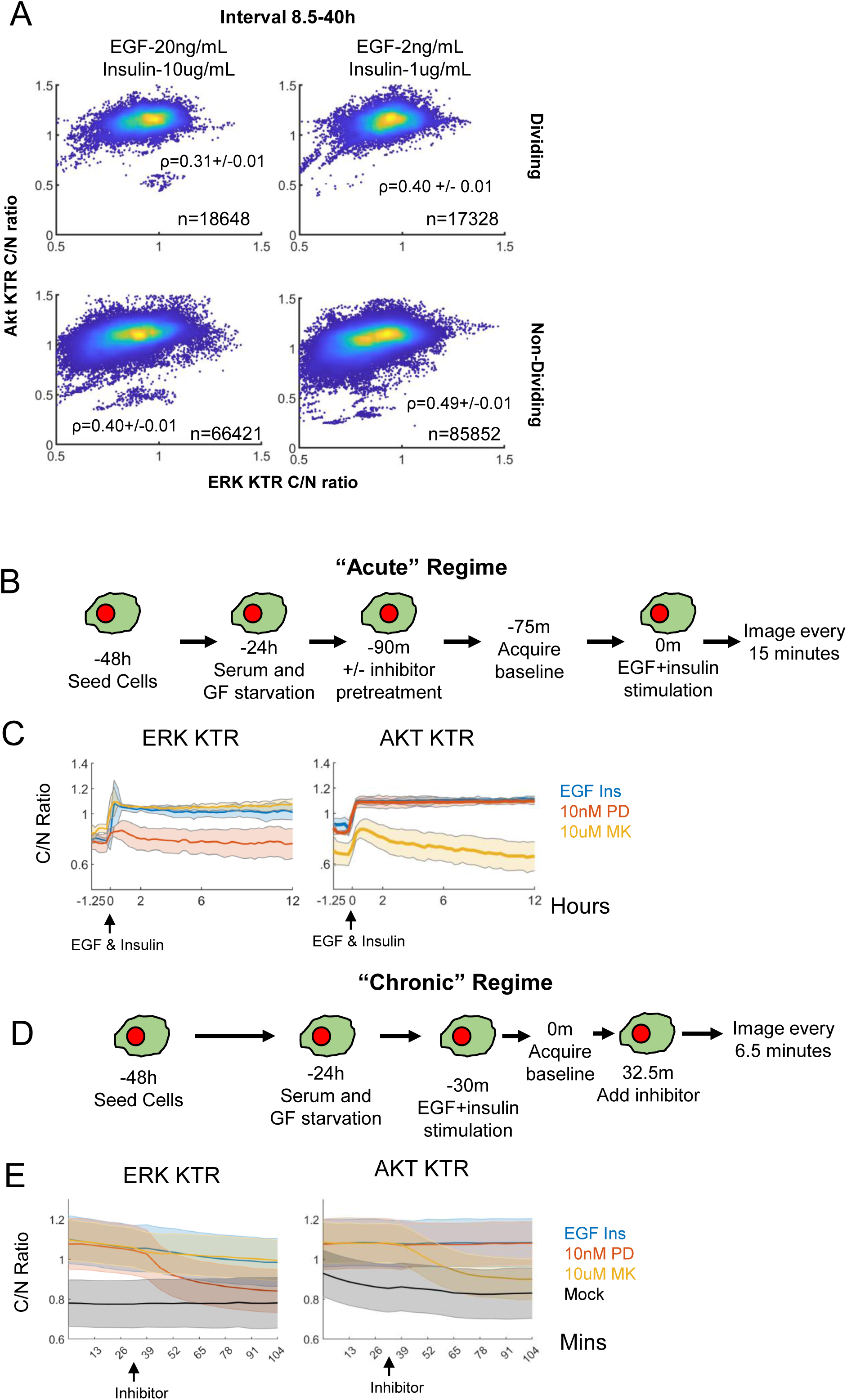
Investigating properties of the ERK and Akt network. (**A**) Single cell ERK and Akt activity plotted across all time points within the 8.5-40 hour interval for dividing and non-dividing cells. These cells expressed both the ERK and Akt KTR. Pearson correlation coefficient, along with the number of cell-time datapoint combinations are indicated. Uncertainty in the correlation coefficients is calculated as described in Methods. (**B**) Cell treatment workflow for network reconstruction in the “acute” regime. Single ERK or Akt KTR expressing MCF10A cells were seeded, allowed to attach overnight, and then serum and growth factor starved. Following starvation, inhibitor was added (PD: PD0325091; MK: MK2206), baseline images were acquired, cells were treated with EGF and insulin, and then imaged every 15 minutes for 12 hours. Images were quantified using the analysis pipeline described in the methods. (**C**) Quantified ERK and Akt activity dynamics in the acute regime. Solid lines are population median (per time point), and shaded areas denote the standard deviation. (**D**) Cell treatment workflow for network reconstruction in the “chronic” regime. Dual ERK and Akt KTR expressing MCF10A cells were seeded, allowed to attach overnight, and then serum and growth factor starved. Following starvation, EGF and insulin were added, baseline images were acquired, cells were treated inhibitor, and then imaged every 6.5 minutes for the remaining ∼hour. Images were quantified using the analysis pipeline described in the methods. (**E**) Quantified ERK and Akt activity dynamics in the chronic regime. Solid lines are population median (per time point), and shaded areas denote the standard deviation.

To reconstruct the ERK and Akt network in MCF10A cells, we implemented recent theory from our lab that specifies a sufficient experimental design for such tasks, based on perturbation time course data^81^. Specifically, for this 2-node network, three time course experiments should be done: response of ERK and Akt activity to EGF and Insulin co-treatment with (i) no inhibitor; (ii) an ERK pathway inhibitor; and (iii) an Akt pathway inhibitor. Additionally, we wanted to understand whether the network would be different in the acute phase of growth factor treatment from a serum starved state vs. the “chronic” condition where ERK and Akt activities are steady over time, particularly because these time regimes seem to have different biological information encoded for cell division decisions.

In the acute regime (Fig. 3B), MCF10A cells expressing either ERK or Akt KTR were seeded, serum and growth factor starved for 24 hours, and then pretreated with either a MEK (PD0325901) or an Akt (MK2206) inhibitor for 30 minutes. The concentrations utilized were determined via titration experiments shown to ensure minimum possible doses were being used (Fig. S1, S2, S4). Following drug treatment, baseline KTR activity was acquired every 15 minutes for 1 hour. Then, we treated cells with EGF and insulin and imaged. Single cell traces (thin) and population median activity (bold) were calculated for each condition, showing robust ERK and Akt activation (Fig. 3C). MEK inhibition ablates ERK activation and has a negligible effect on Akt activation (Fig. 3C). Akt inhibition ablates Akt activation and has a negligible effect on ERK activation (Fig. 3C). Although the Akt KTR may reflect kinase activity other than Akt in this acute stimulus regime, the fact that complete inhibition of the ERK pathway has negligible impact on the Akt KTR readout means that ERK does not impact Akt or the others. These results show that in the acute stimulus regime, ERK and Akt exhibit negligible crosstalk after treatment with EGF and insulin.

In the chronic regime, cells were pretreated with either EGF and insulin for 30 minutes followed by 30 minutes of baseline acquisition, leading to robust ERK and Akt activation (Fig. 3D-E, Replicates in S5). Akt inhibition reduces Akt activity, as expected, but negligibly affects ERK activity. MEK inhibition reduces ERK activity, as expected, but does not appreciably affect Akt activity. These conclusions are also consistent when a second set of MEK and Akt inhibitors are used (Fig. S4). Therefore, in the chronic regime ERK and Akt also do not exhibit appreciable cross pathway interactions after EGF and insulin co-treatment. We conclude it is unlikely that crosstalk interactions account for correlations that change in dividing vs. non-dividing cells.

## Discussion

Binary single-cell responses, like division, to perturbations such as growth factor and drug treatments, are almost universally heterogeneous even in clonally derived populations. However, predictive biochemical features, present either before the perturbation, or from dynamics following the perturbation, are seldom known. The ability to predict such binary responses would not only reflect a deep and fundamental understanding of the systems governing important cellular responses, but also have significant translational applications such as antibiotic resistance, tissue engineering, and anticancer therapy, where the fates of single cells can be of great importance. Here, we investigated growth-factor induced cell division fates in the well-studied, non-transformed mammalian epithelial cell line MCF10A, and how they may be predicted by the dynamics of two central signaling pathways, PI-3K/Akt and Ras/ERK. Answering such questions requires single-cell, non-destructive analysis of biochemical features, in this case ERK and Akt activities, that are paired to the eventual cell division outcome. They also must be carried out in a high-throughput manner to observe enough events to make statistically-supported conclusions. After setting up this experimental system and understanding its ranges of validity, we learned that (i) ERK and Akt activities are higher in the 8.5-40 hour window after growth factor treatment in cells that divide, suggesting underappreciated roles post-restriction point, into S and G2 phases; (ii) median activities in this time window predict single cell division outcome with AUC=0.76 with ERK dynamics alone giving AUC=0.74, suggesting Akt dynamics add little to the decision driving division in this context; (iii) metrics of central tendency are the most predictive features; (iv) ERK and Akt activities are less correlated in cells that divide; and (v) ERK and Akt do not exhibit crosstalk in this system, so division-related correlation is unlikely related to crosstalk.

We have performed these studies in the MCF10A cell line, a well-established model for non-transformed epithelial cells. An obvious next question is how the relationships between ERK, Akt and cell division found here translate to different cell lines, and transformation contexts. Many other cell lines are cancer-derived and genetically unstable, and/or contain multiple alterations to the systems that control cell cycle progression and division. A firm understanding of how ERK and Akt systems control the cell cycle in a system such as MCF10A is an important foundation for subsequently understanding how it may be altered in other cell lines, and also across different microenvironmental contexts, such as confluent settings. Indeed, there is a growing body of work that focuses on answering fundamental cell biological questions using studies on the MCF10A system alone (e.g. ^82^). It is appealing to consider MCF10A as an emerging model system for mammalian epithelial cells.

Nearly all the cells we observed had relatively simple dynamics for ERK and Akt activity, a rise then a somewhat constant higher than baseline steady-state. Other recent single cell studies have reported pulsatile ERK dynamics^1,49,83,84^. Some of this may be related to differences in growth factor concentration, the reporters used, being FRET-based ^85^ or translocation based ^70^. No live-cell imaging probe is perfect and of course has its drawbacks, some of which may be related to off-target responses, which may partly explain our Akt activity data in the “acute” phase first following growth factor treatment. For example, kinases other than Akt may recognize and phosphorylate the FOXO1-based Akt KTR docking site ^86–90^. EGF and insulin stimulation may also promote activation of such kinases including PLK1 ^91^, SGK and PKA^92^. Another aspect may have to do with cell-cell contact and density. In our study, cells were seeded at low density and serum/growth factor-starved prior to analysis, whereas pulsatile signaling was reported in high density environments in asynchronously cycling settings^1,84^. Yet others have found pulsatile dynamics can induce different sets of genes as compared to sustained dynamics^46^. However, phenotypic consequences, at least in terms of cell proliferation still seem to be related to simple time-integrated signaling dynamics^1,50^, similar to what we found here.

ERK and Akt activity dynamics are only a subset of the potentially important variations that drive phenotypic variability in cell division responses, as shown by the AUC=0.76 that was obtained. ERK dynamics account for nearly all this predictive power. This reinforces Akt activity as perhaps more relevant for cell maintenance and health, and more as a “checkpoint” for division but not a significant driver, at least in the studied system. As noted above, cell contacts and density are important. Such phenomena may potentially be controlled through micropatterning experiments, where cell shape and placement can be carefully controlled^93,94^. Cell “state”, corresponding to different epigenetic and/or metabolic states of cells prior to the experiment, has been reasonably well documented ubiquitously, and can contribute to variability, although is difficult to assess in the “track and follow” manner that can be done with live-cell kinase reporters. Metabolic or organelle abundance variability may also contribute^95,96^. Of course, there are other pathways and biochemical correlates that are likely important, such as a balance between p53 and p21 and/or CDK2 activity^54,67,97^. Given the multitude of fluorescent proteins, and improvements in cell tracking from non-labeled bright field images^98,99^, one may be able to measure more important biochemical readouts simultaneously for such purposes. There are also multiple checkpoints between growth factor treatment and cell division, such as the restriction point, and DNA damage checkpoints, that may contribute. Monitoring division with probes like the Fucci system that gives readouts of each cell cycle phase may help explore such phenomena^100^.

An interesting aspect of our study was the surprising larger differences between dividing and non-dividing cells in the time period that corresponds to S/G2 phases of the cell cycle, as opposed to pre-S-phase. The roles of growth factor signaling through ERK and Akt pathways historically focused on passing the restriction point into S-phase^101^. Thus, our results suggest potential functional roles for ERK and Akt beyond this canonical understanding. Indeed, a recent study found time-integrated ERK activity in a mother cell’s G2 phase influenced the cell cycle progression in the subsequently daughter cells^50^. The mechanisms that may be driving such functional roles are a potentially interesting area of future study.

We also studied the ERK and Akt activity network, since we found that ERK and Akt activity are less correlated with each other in dividing cells compared to non-dividing cells. We found that the observed differences in correlation are likely not a result of network topology as ERK and Akt do not appreciably interact. This lack of interaction is surprising given that some prior studies describe these pathway as exhibiting cross pathway interactions ^23,69,75^, albeit in other cell lines and in response to other growth factors. However, other studies in MCF10A cells across a panel of different growth factors show that ERK and Akt are insular, and do not interact ^69^. Similar to MCF10A cells, 32D-EpoR; BaF3-EpoR; CFU-E cell lines exhibit minimal ERK and Akt cross pathway interaction under erythropoietin stimulation, a growth factor that activates both ERK and Akt ^80^. These studies reveal that in non-interacting pathways, differences in protein expression influence the flow of erythropoietin signaling ^80^. Therefore, in our model system, it is possible that the observed differences in ERK and Akt correlation may arise from differences in protein expression across dividing and non-dividing cells. It may also be that differences in upstream signaling capacity to ERK and Akt may be related. Characterizing the differences in protein expression level in single cells, and following cell signaling and cell division can provide insight; but this becomes a quite challenging experiment given the number of probes to be measured simultaneously.

In conclusion, we have studied the relationship between ERK activity dynamics, Akt activity dynamics, and cell division, and found that simple measures of central tendency of these dynamics in a time coinciding with S/G2 phase are most predictive of cell division in single cells. This implies unappreciated roles for ERK and Akt beyond the canonical restriction point. ERK accounts for much of this predictive capacity, suggesting Akt contributes little to the decision to divide in this context. Yet, AUC=0.76 is far from perfect so it is clear other biochemical pathways are important factors for predicting single cell division events. ERK and Akt do not interact with one another in the studied contexts, despite the fact that their activities are more correlated in cells that do not divide. These studies conducted in the non-transformed context provide a foundation to explore how cell transformation through oncogenic and/or loss-of-function mutations shape network topology, signaling dynamics, and cell division outcome in cancer, with the potential to identify and target pathway compensation behaviors that promote cell proliferation and survival ^102^ in the diseased context. In addition, exploring the role of spatial temporal propagation of ERK and Akt signaling in a 3D tissue context, a model system that MCF10A cells are suited for, can provide insight into how these pathways regulate tissue homeostasis and how transformation disrupts this homeostasis.

## Methods

### Cell Culture

MCF10A cells were gifted by Dr. Gordon Mills and cultured in complete sterile filtered (VWR 10040-436) media, consisting of DMEM F12 (Gibco #11330-032) supplemented with 2 mM L-Glutamine (Gibco # 25-005-CI), 20ng/mL EGF (Peprotech AF-100-15), 10ug/ml insulin (Sigma #I-1882), 0.5ug/ml hydrocortisone (Sigma #H-0888), 100ng/ml cholera toxin (Sigma #C-8052) and 5% horse serum (Invitrogen #16050-122). Cells were passaged with 0.25% trypsin (Gibco #25200056) to maintain sub confluency. Cells were maintained at 37°C, 5% CO_2_. Starvation media and imaging media is phenol red free DMEM F12 (Fisher #11039-021) supplemented with 100ng/ml cholera toxin.

HEK293T cells were gifted by the Dr. Dominguez and Dr. Pappapetrou labs and cultured in complete sterile filtered (VWR 10040-436) media, consisting of DMEM (Gibco #11965118) supplemented with 2 mM L-Glutamine (Gibco #25-005-CI) and 10% heat inactivated fetal bovine serum (Gibco #16140071). Cells were passaged with 0.05% trypsin (Gibco #25300054) to maintain sub confluency.

All inhibitors used for KTR validation were formulated as 10mM stock solutions in DMSO (Sigma Aldrich D2650-5X0ML) and sterile filtered with a 0.22-micron syringe filter. PD0325901 (MEK inhibitor 1) was purchased from Sigma Aldrich (PZ0162-5MG). Trametinib (S2673) (MEK inhibitor 2) and Ipatasertib (S2808) (Akt inhibitor 2) were purchased from Selleck Chemicals. MK2206 (#CT-MK2206) (Akt inhibitor 1) was purchased from Chemietek.

### Imaging

All live cell imaging experiments were acquired using the InCell 2200 microscope (GE Healthcare) under environmental control (37°C, 5% CO_2_.) with a Nikon 20X/0.75, Plan Apo, CFI/60 objective. For KTR imaging the following filter sets were utilized: FITC (Excitation: 475/28nm Emission: 511/23nm) (ERK-mClover, Akt-mClover KTR); Cy3 (Excitation: 542/27nm Emission:597/45nm) (H2B-mRuby2, mCherry-NLS); Cy5 (Excitation: 632/22nm Emission: 679/34nm) (ERK-iRFP); Brightfield.

KTR-expressing MCF10A cell lines (see below) were seeded in separate rows of a 96 well plate (Corning #3603) at 5000 cells / well and treated as described. After growth factor and serum starvation, starvation media was aspirated, cells were washed with PBS and 100 uL imaging media was then placed in the wells. Following baseline imaging, cells were treated as indicated by adding 100 uL of 2X solutions in imaging media. Acquired images were processed as described *Computational Image Analysis*.

### Flow Cytometry

EdU flow cytometry assays were performed using the Molecular Probes Click-iT Plus EdU flow cytometry assay kit (C10633 molecular probes). MCF10A cells were seeded in 6 well plates (Corning 353046) at a density of 127 cells/mm^2^ in complete DMEM F12 media. The following day, we serum and serum and growth factor starved cells in DMEM F12 media supplemented with 100ng/mL cholera toxin for 24 hours. Following starvation, cells were pretreated with a final concentration of 100nM of MEK inhibitor 1 (PD0325901) or 10uM of Akt inhibitor 1 (MK2206) or DMSO control for 30 minutes. Cells that did not receive inhibitor pretreatment were treated with either MEK inhibitor 1 or Akt inhibitor 1 2, 4, 8, or 12 hours post EGF and insulin addition (Final growth factor concentration: 20ng/mL EGF, 10ug/mL insulin). 22 hours post growth factor addition, a final concentration of 10 uM of EdU was added to each well and incubated for two hours. Two hours post EdU addition, cells were washed with PBS and lifted with 0.25% Trypsin for 10 minutes. Trypsin was neutralized with complete DMEM F12 media. Cells were pelleted at 100xG for 5 minutes, resuspended in 100uL of PBS, and processed as recommended by the manufacturer’s protocol (Molecular Probes Click-iT Plus EdU flow cytometry assay kit). During the last 5 minutes of permeabilization, 100uL of diluted 1ug/mL Hoechst 33342 (Thermo Fisher H3570) stain was added. Cells were then washed with a 1 % (g/100mL) bovine serum albumin PBS solution and pelleted at 100g. Cells were resuspended in permeabilization buffer and stained with EdU Click-iT reaction cocktail for 30 minutes at room temperature protected from light. Following EdU Click-iT labeling, cells were washed and resuspended in permeabilization buffer and analyzed using the BD Canto II flow cytometer configured with the following laser lines: excitation 640nm, emission filter 660/20, excitation 405nm, and emission filter 450/50. Data were gated and processed using FCS Express (Denovo Software).

### Cloning

Akt KTR was modified from the transposase transfection system PSBbi-FoxO1_1R_10A_3D vector (Addgene # 106278)^48^ for lentiviral production (Fig. S6A). The lentiviral expression vector was developed using overlap PCR from fragments generated from the source vector PSBbi-FoxO1_1R_10A_3D: SV40NLS-mCherry-P2A and Gly-FT2DDD-KTR-mClover. Primer sequences are shown in Table S1 and were designed in SnapGene and ordered from Sigma Aldrich. Fragments for lentiviral vector construction were generated via PCR using Q5 polymerase (NEB M0491S) and primers specific to SV40NLS-mCherry-P2A and Gly-FT2DDD-KTR-mClover (Table S1) regions of PSBbi-FoxO1_1R_10A_3D (Fig. S6, Fragment 1,2). Fragments were gel purified using the NEB Monarch gel extraction kit (NEB T1020S). Following gel extraction, a 10 cycle PCR reaction was performed using Q5 polymerase and equimolar SV40NLS-mCherry-P2A and Gly-FT2DDD-KTR-mClover fragments using an annealing temperature of 72°C. 5 uL of the product was amplified using end primers (F: SV40NLS-mCherry-P2A, R: Gly-FT2DDD-KTR-mClover, T_a_=69°C, Table S1). Gateway ATTB sites were inserted at flanking ends using PCR and the ATTB primers (Table S1). The Gateway cloning compatible fragment was inserted into donor vector pDONR221 (Invitrogen™ 12536017) using BP Clonase II (Thermo Fisher# 11789020). High Efficiency NEB-5-alpha Competent E. coli (NEB C2987I) were transformed with pDONR221 containing the Akt KTR. Transformants were miniprepped with the PureYield™ Plasmid Miniprep System (Promega A1223) and Sanger sequenced verified with GeneWiz. Akt KTR expression vector was generated by performing a LR reaction using pDONR 221-Akt, LR Clonase II (Thermofisher #11791020) and the lentiviral expression vector pLenti CMV Hygro DEST (Addgene #17454) generating the final product, a bi-cistronic hygromycin selectable lentiviral expression vector. The product was transformed into NEB® 5-alpha Competent E. coli. Transformations with the correct sequence were maxiprepped with PureYield™ Plasmid Maxiprep System (Promega # A2392) and utilized for lentiviral production.

We exchanged the antibiotic selectable marker on the lentiviral expression vector H2B-mRuby2 (Addgene #90236) from hygromycin to puromycin. Specifically, we transferred pLentiPGK Hygro DEST H2B-mRuby2 into pLentiCMV puromycin DEST (Addgene #17452) using BP Clonase II followed by LR Clonase II generating pLentiCMV puromycin DEST H2B-mRuby2 (H2B-mRuby2).

### Lentiviral Production

The lentiviral constructs for each cell line are shown in Fig. S6B. Lentiviral particles were generated by transfecting 5 million HEK293T cells seeded in a T75 flask and allowed to attach overnight (Corning^®^ T-75 flasks catalog #430641) using the TransIT-293 transfection reagent (Mirus Bio MIR2704) along with expression vector ERK KTR (mClover or iRFP), H2B-mRuby2, or Akt KTR along with packaging plasmid pPAX (Addgene #12260), and envelope protein pCMV-VSV-G (Addgene #8454) according to the manufacturer’s instructions. Two days post transfection, supernatant from was collected and concentrated using Amicon Ultra-15 100 kD centrifugation filters (Millipore #UFC910008). The concentrated lentiviral supernatant was aliquoted and stored at -80°C.

### Lentiviral Transduction

100,000 MCF10A cells were transduced in suspension in a 6 well plate (Corning 353046) containing complete DMEM F12 medium along with 100uL lentiviral supernatant. Two days later, expression was validated by fluorescence imaging. ERK KTR MCF10A cell lines (ERK KTR-mClover Hygro, H2B-mRuby2 Puro) were selected in complete DMEM F12 media supplemented with hygromycin (35ug/mL) and puromycin (2ug/ml). Akt KTR expressing MCF10A cell lines (SV40nls-mCherry-Akt KTR-mClover Hygro) were selected with DMEM F12 media containing hygromycin (35ug/ml). Cells were passaged every two to three days in selection media for about two weeks. Following selection, cells were expanded in complete DMEM F12 maintenance media containing either both hygromycin (1.5ug/ml) and puromycin (0.1ug/ml) (ERK KTR-mClover expressing cells) or hygromycin (1.5ug/ml) (Akt-KTR-mClover expressing cells). Live-cell imaging was conducted in the absence of selection antibiotics. Dual reporter expressing cells (ERK KTR-iRFP, Akt-KTR-mClover) lines were not selected, as the ERK KTR iRFP (Addgene #59150) lentiviral expression vector does not confer antibiotic resistance. For these, ERK KTR iRFP virus was added to cells for 24 hours, and then subcultured as above prior to live-cell imaging analysis.

### Computational Image Analysis

While many image analysis tools exist ^103–105^, each application still requires much novel development tuned to the problem at hand. We developed an automated image analysis pipeline using both iLastik ^106^ and CellProfiler ^105^ software packages, along with MATLAB scripts (Fig. S2C). It is available at the Birtwistle Lab github repository (github.com/birtwistlelab/Predicting-Individual-Cell-Division-Events-from-Single-Cell-ERK-and-Akt-Dynamics), which includes some dockerized scripts. The analysis pipeline consists of (1) cell nuclei and cytoplasmic segmentation, (2) quantification of KTR fluorescence in both nuclei and cytoplasmic compartments, (3) tracking single cells across a time series, and (4) automatic detection of cell division.

1. Prior to segmentation images were flatfield corrected and background subtracted using CellProfiler. Images of nuclear localized fluorescent protein H2B-mRuby2 (ERK KTR) and NLS-mCherry (Akt KTR) were input into iLastik. Nuclei were identified using a series of features-object intensity, edge detection and texture.
2. The binary mask outputs from iLastik were input into CellProfiler to create a perinuclear ring known as the ‘Cytoring’^107^ which extends 10 pixels from the binary nuclei mask and into the cytoplasm (Fig S2C). Calculating the cytoplasmic to nuclear KTR fluorescence ratio provides the relative activity of the pathway of interest for that particular cell at that time point.
3. Segmented nuclei identified with iLastik were tracked using CellProfiler’s ***TrackObjects*** module ^104,105,108^ and Follow Neighbors^108^. Each identified nucleus was assigned a numerical ID, which corresponds to the same cell across each timepoint. We filter tracks that are shorter than the duration of the time course to prevent quantification of cells that were transiently tracked.
4. Cell division was detected using a feature of cytoplasmic to nuclear KTR fluorescence (C/N ratio) that is unique to dividing cells. As cells divide, there is a change in morphology resulting in a rapid decrease in C/N ratio (Fig. S7). MATLAB’s *findpeaks* function was used to detect when this steep decrease occurs. We then truncated the time series 5 timepoints before the identified peak, which is attributed to actual kinase activity.

The CellProfiler pipeline exports CSV files first preprocessed in Microsoft Excel then analyzed in MATLAB. First, the csv are input into **batchreader.m,** which generates a cell array of tables containing each cell’s measured parameters. The data is then input into the script **ktrTablePlotter.m,** which plots KTR dynamics. Cell division events are detected using the script **Div_detection.m.** Division events were confirmed by directly observing nuclear fission via H2B-mRuby2 or NLS-mCherry fluorescence. Cells were separated by division status, and KTR dynamics were plotted for each class.

### Statistics, Classification, and Visualization

Rank-sum tests were used to calculate p-values for differences between dividing and non-dividing cells. For single reporter expressing lines the MATLAB function *fitglm* was used with median ERK or Akt activity within the 8.5-40 hour interval as the predictor class. ROC curves were generated using the MATLAB function *perfcurve*. For dual ERK and Akt KTR expressing MCF10A cell lines, 150 randomly selected dividing and non-dividing cells were used to develop a linear SVM classifier using the MATLAB function *fitcsvm*. The MATLAB function *resubPredict* was used to calculate the SVM classification performance between the two classes.

NotBoxPlot was retrieved from MATLAB Central File Exchange (Rob Campbell, 2021). Scatter plots of single cell ERK and Akt activity across all timepoints within the 8.5-40 hour interval were generated using the MATLAB Central File Exchange script *Scatplot.m* - Alex Sanchez (2020). To assess statistical significance of the correlation coefficient, the mean (µ) and covariance (σ) between ERK and Akt were calculated across all biological replicates and used to sample matched numbers of data points from random multivariate normal distributions for dividing cells and non-dividing cells. This was repeated 1000 times to define the range of correlation coefficients between the 5^th^ and 95^th^ percentiles, which was reported and rounded up to the nearest 0.01.

The hctsa package^72,73^ was installed according to the published documentation. We initialized our data set from data acquired under high concentrations of EGF and insulin stimulation using the custom MATLAB script *hctsa_prep_dualrep.m*. The number of cells that were input into hctsa were chosen based on the smallest population of either dividing /non-dividing cells. A random permutation was then used to choose an equal number of the largest population of cells to input into hctsa. The data set was initialized using the hctsa’s *TS_init* command, followed by *TS_compute* command. The processed datasets were labeled for hctsa format as 1-dividing or 0-non-dividing using

### TS_LabelGroups

The top predictive features were identified using the raw computed hctsa values and the function *TS_TopFeatures* for both ERK and Akt time course data.

---

**Figure S1.**
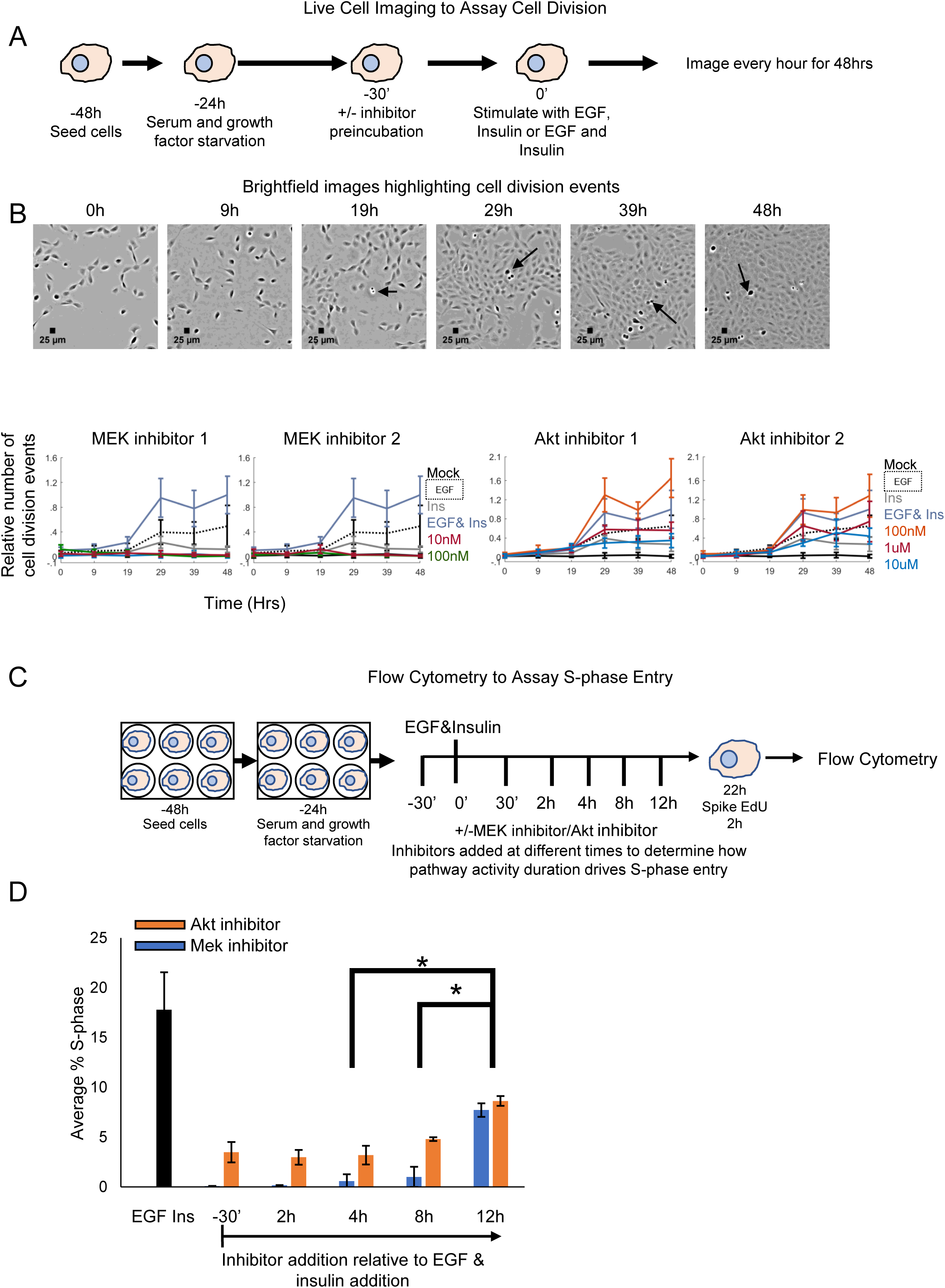
Roles of ERK and Akt activity in cell cycle progression in MCF10A cells. (**A**) Live cell imaging pipeline for quantifying cell division events in MCF10A cells under EGF, insulin, and EGF & insulin stimulation. MCF10A cells were seeded, serum and growth factor starved as described in the methods. Following starvation, cells were pre-incubated with or without MEK inhibitors 1,2 (PD0325901, Trametinib) or Akt inhibitors1,2 (MK2206, Ipatasertib) for 30 minutes. Following inhibitor preincubation, growth factors EGF (20ng/mL), insulin (10ug/mL), or EGF (20ng/mL) & insulin (10ug/mL) were added. Cells were imaged under brightfield every hour for 48 hours. (**B**) Top: Representative brightfield images of cells under EGF and insulin stimulation. Arrows point to representative cell division events. The number of cell division events at selected time points were counted in this manner. Bottom: The relative number of cell division events were calculated by summing the number of observed cell division events across each field of view per condition divided by the total number of observed division events under EGF and insulin stimulation. Error bars represent the standard deviation of normalized cell counts per field per condition. Insulin induces essentially no cell division, but when in combination with EGF, has a more than additive effect. Both ERK and Akt activities appear essential for cell division in this context. (**C**) Experimental outline for relating ERK and Akt dynamics in driving S-phase entry in MCF10A cells. MCF10A cells were seeded, serum and growth factor starved as described in the methods. Post starvation, cells were treated with MEK or Akt inhibitors 1 (PD-10nM and MK-10 uM, according to minimum effective concentrations above) at the times indicated relative to EGF and insulin addition. 22 hours post growth factor addition, EdU was spiked in for 2 hours. (**D**) The average percentage of the cell population in S-phase for each of the conditions shown. A single tailed Student’s t-Test was used to calculate the significance between the average percentage of cells entering S-phase between 4 & 12 hours and 8 & 12 hours at the 95% confidence interval (p< 0.00045). Error bars are from biological triplicates. These results are consistent with the model that time integrated ERK and Akt activities for at least 8-12 hours are necessary for S-phase entry.

**Figure S2.**
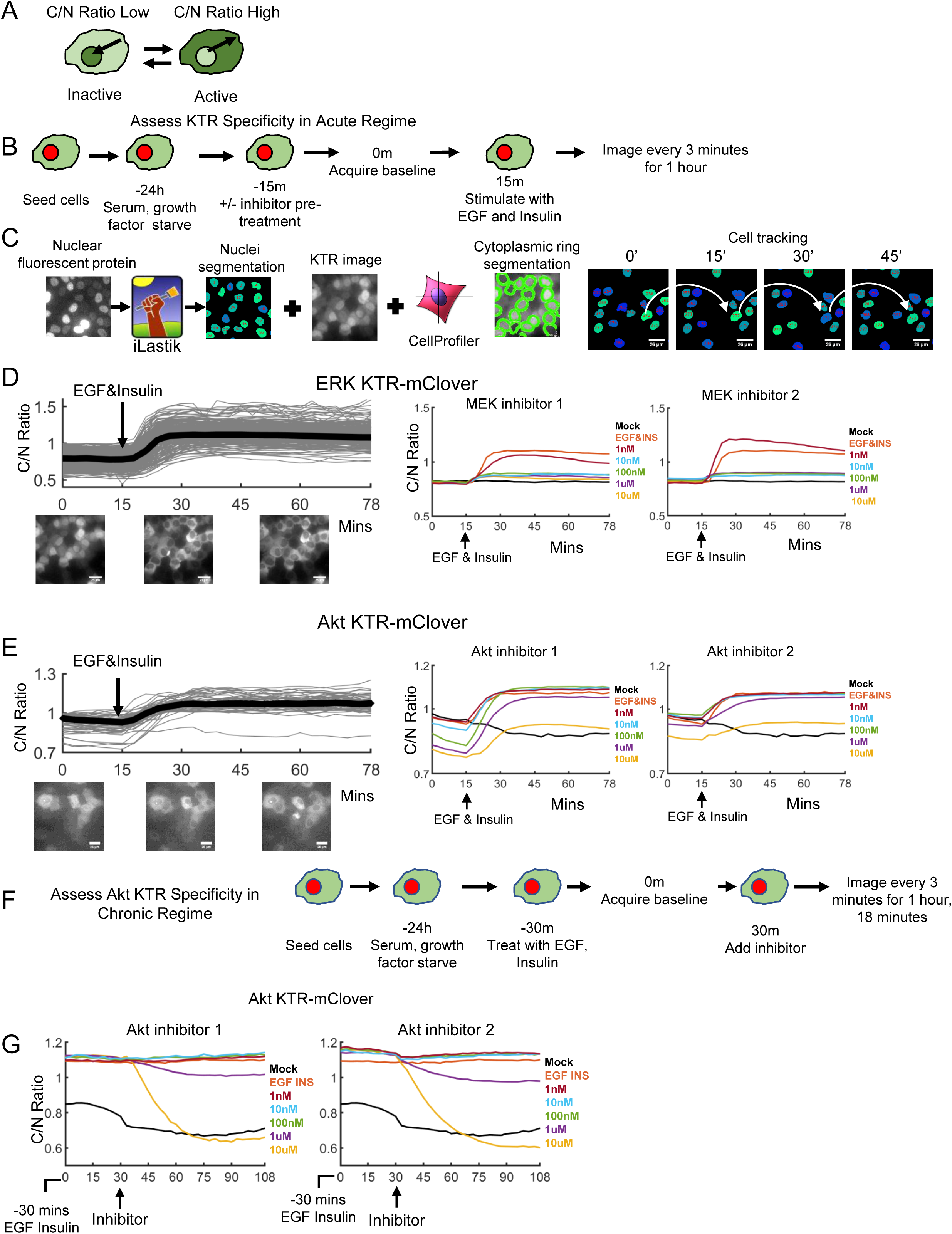
Live cell imaging assays to establish regimes of validity for kinase translocation reporters. (**A**) Cartoon representation of kinase translocation reporter in the inactive (nuclear) and activated (cytoplasmic) state. (**B**) Acute stimulus kinase translocation reporter validation pipeline. (**C**) Steps in the computational image analysis pipeline for nuclei, cytoplasmic identification, cell tracking and quantification of reported KTR dynamics. (**D**) Single cell traces (gray thin lines) of quantified ERK KTR (C/N Ratio) under EGF (20 ng/mL) and insulin (10 ug/mL) stimulation. Population median (per time point) response is shown in thick black. Representative images of cells expressing ERK KTR are shown below each time point: 0, 30, and 60 minutes. Prior to EGF addition, ERK KTR is mainly nuclear localized (0-15 minutes). By 15 minutes post EGF and insulin addition, ERK KTR is cytoplasmic localized, which is reflected by the increase in the C/N ratio. The panel to the right shows the population median C/N traces for each inhibitor dose condition. KTR activity is ablated with ∼10 nM of both MEK inhibitors. MEK inhibitors 1 and 2 are PD0325901 and Trametinib, respectively. (**E**) Single cell traces (gray thin lines) of Akt KTR (C/N Ratio) under EGF (20 ng/mL) and insulin (10 ug/mL) stimulation. Population median (per time point) response is shown in thick black. Prior to EGF and Insulin stimulation, Akt KTR exhibits slightly elevated C/N Ratio, indicative of some basal activity. By 15 minutes post growth factor stimulation, Akt KTR is cytoplasmic localized as shown by C/N Ratio values. Shown to the right are the population median traces for each inhibitor dose condition. Even at 10 uM of each Akt inhibitor, there is significant Akt KTR activity, indicating that at least in the first ∼1 hour post-EGF and insulin treatment, the Akt KTR activity likely reports on other kinases. Akt inhibitors 1 and 2 are MK2206 and Ipatasertib, respectively. (**F**) Live cell imaging pipeline to investigate Akt KTR validity under chronic EGF and insulin stimulation. (**G**) Population median Akt KTR dynamics for each inhibitor dose condition. After 1 hour of EGF and insulin treatment, 10 uM of either Akt inhibitor completely ablates Akt KTR reporter activity.

**Figure S3.**
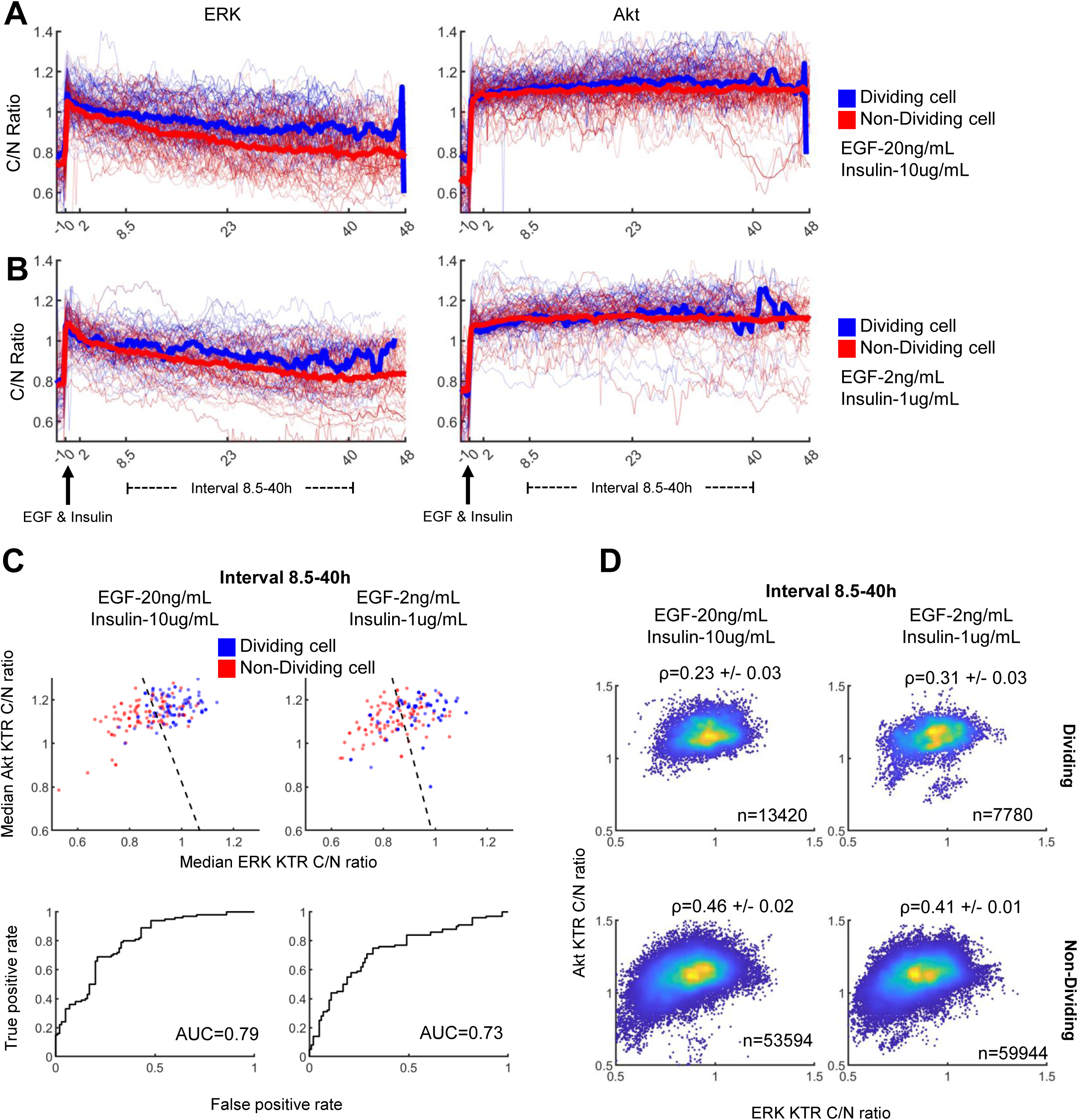
Biological replicates for data presented in Figures 2 and 3. Please see those legends for details.

**Figure S4.**
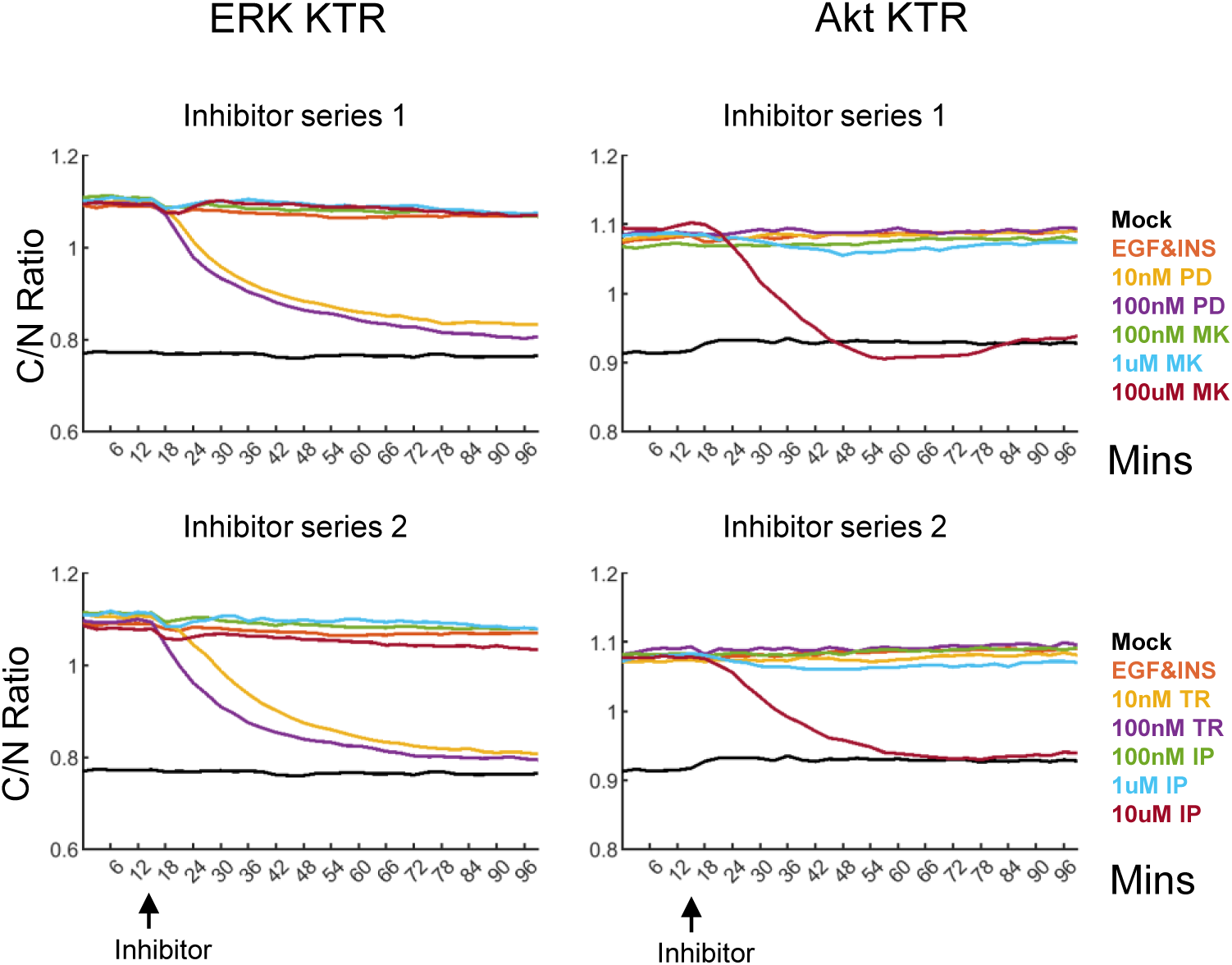
Biological replicates for KTR range of validity and network reconstruction in the chronic setting using single reporter cell lines. MCF10A cells expressing either ERK or Akt KTR were pre-incubated with EGF and insulin for 90 minutes followed by baseline KTR acquisition. Following baseline, either a MEK or an Akt inhibitor was added. Population median KTR activity calculated across cells for each timepoint are shown for both inhibitor 1 and 2. MEK inhibitors 1,2 are PD0325901 and Trametinib and Akt inhibitors 1 and 2 are MK2206 and Ipatasertib.

**Figure S5.**
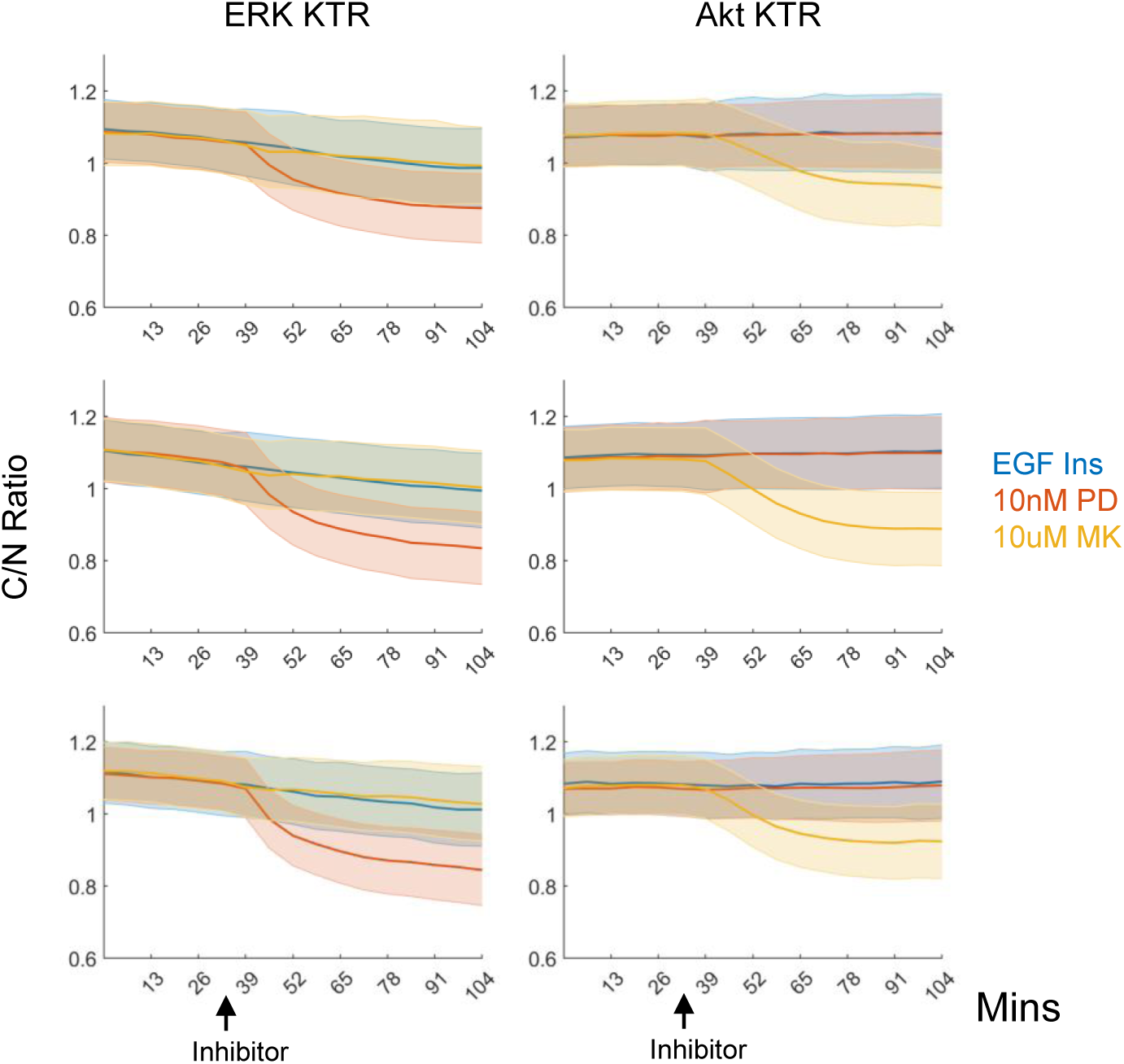
Biological replicates for chronic regime network reconstruction. Thick lines are population median (per time point), and shaded regions +/- standard deviations across single cell responses. Please see Figure 3 for more details.

**Figure S6.**
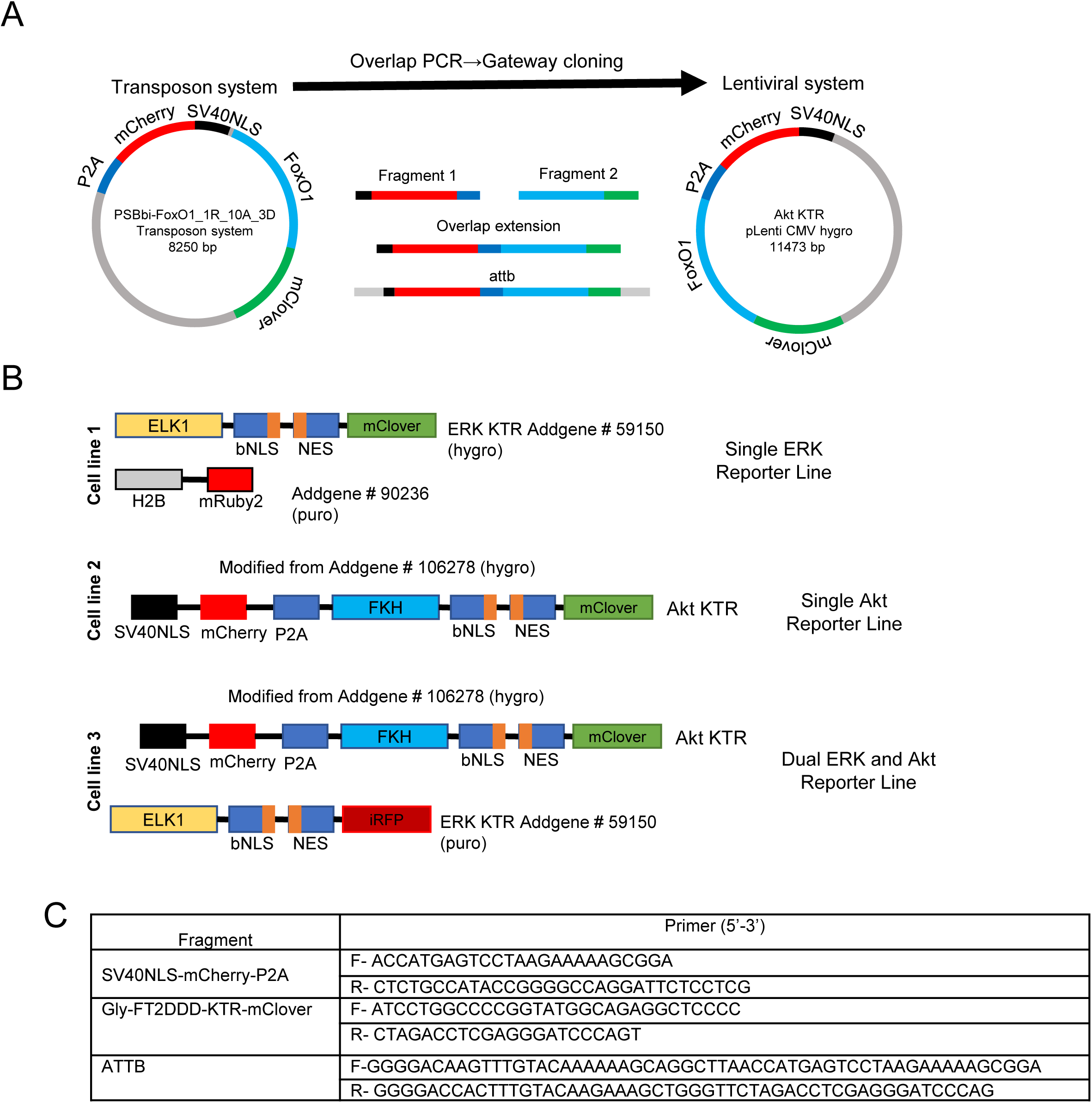
KTR construction and cell lines. (**A**) Converting the Akt KTR from the transposon backbone to the lentiviral backbone. (**B**) Cartoon representation of constructs used to generate KTR expressing MCF10A cell lines. (**C**) PCR primers used for the different vector construction steps.

**Figure S7.**
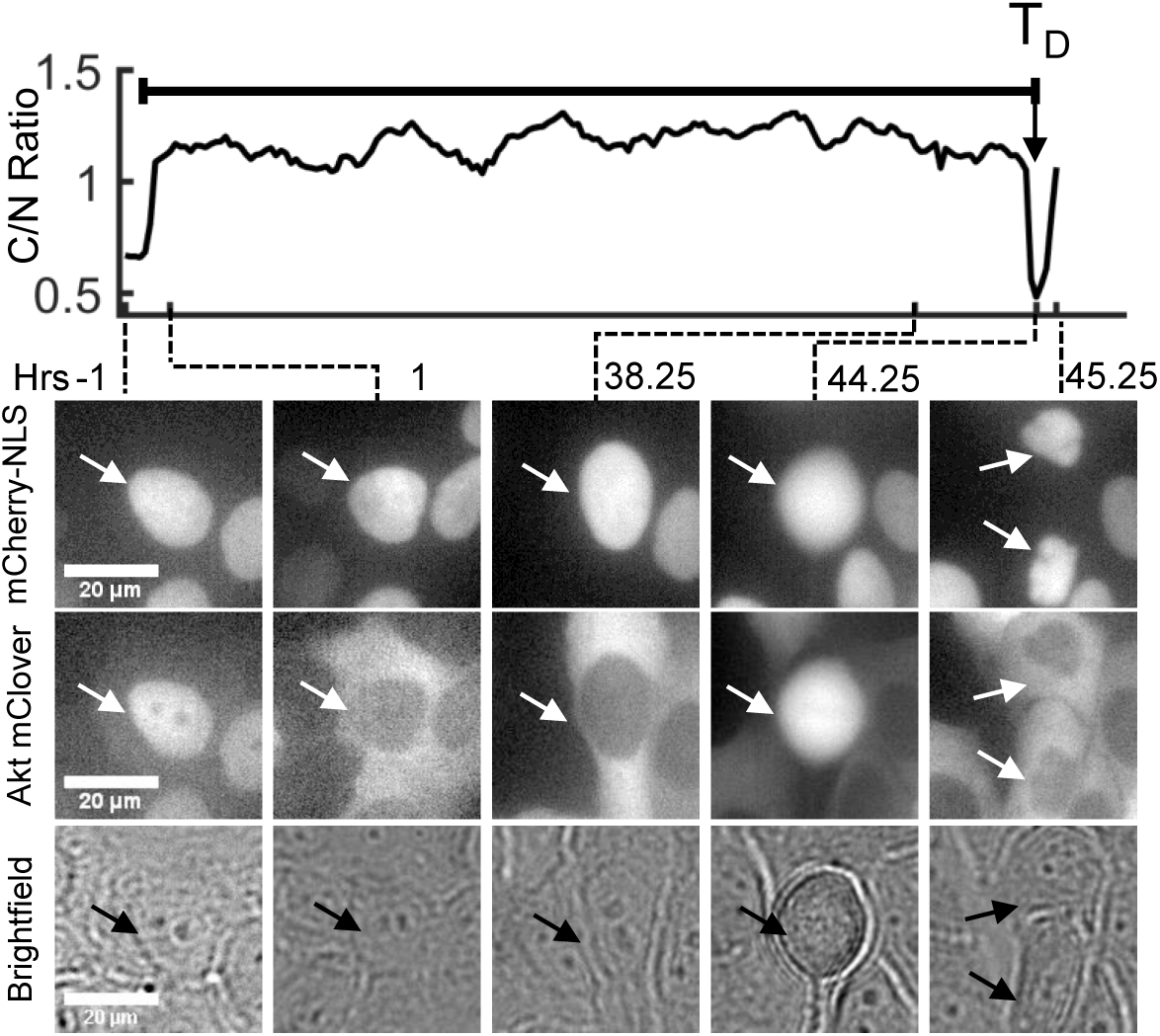
Identification of cell division events. A representative time course and selected images from an Akt KTR expressing MCF10A cell that divides in response to EGF and insulin treatment is highlighted. The nuclear marker is mCherry-NLS. T_D_ denotes the time of division which is determined by the rapid decrease in C/N ratio.

